# Two distinct mechanisms lead to either oocyte or spermatocyte decrease in *C. elegans* after whole developmental exposure to γ-rays

**DOI:** 10.1101/2023.04.26.538382

**Authors:** E. Dufourcq Sekatcheff, C. Godon, A. Bailly, L. Quevarec, V. Camilleri, S. Galas, S. Frelon

## Abstract

Wildlife is subject to various sources of pollution, including ionizing radiation. Adverse effects can impact organisms’ survival, growth, or reproduction, later affecting population dynamics. In invertebrates, reproduction, which directly impacts population dynamics, has been found to be the most radiosensitive endpoint. Understanding the underlying molecular pathways inducing this reproduction decrease can help to comprehend species-specific differences in radiosensitivity. In line with previous studies, we used a life stage dependent approach to better understand the molecular determinants of reproduction decrease, especially of gamete decrease, in the hermaphrodite roundworm *Caenorhabditis elegans*. Worms were chronically exposed to 50 mGy·h^−1^ external gamma ionizing radiations throughout different developmental periods (namely embryogenesis, gametogenesis, and full development). Conserved molecular pathways across invertebrates and vertebrates involved in reproduction processes and stress response were analyzed: apoptosis and MAP kinase Ras/ERK (MPK-1). Our results showed that these pathways are life-stage dependant, resulting from an accumulation of damages upon chronic exposure to IR throughout the whole development. The Ras/ERK pathway was found to be activated in our conditions in the pachytene region of the gonad where it induces the apoptotic pathway, but not in the ovulation zone, showing no incidence on oocyte maturation and ovulation. Additionally, no effect on germ cell proliferation was found, meaning that Ras/ERK pathway is probably not involved in this process in our conditions. Finally, a functional analysis of apoptosis revealed that the decrease of ovulation rate is due to DNA-damaged induced apoptosis which does not occur in spermatocytes. Thus, sperm decrease seem to be mediated via another mechanism, probably a decrease in germ cells proliferation speed that needs further investigation to better characterize sex-specific responses to IR exposure.

## INTRODUCTION

Because of the increased number of pollutants that accumulate in the environment every year, there is a growing need to expedite and enhance their screening for toxicity. Risk assessment for ecosystems *per se* is a growing concern, especially since the Convention on Biological Diversity of 1992, all the more in the one health concept cross-connecting animals’, ecosystems’ and Human health (1). This assessment requires both a robust knowledge on the relationship between environmental exposure and induced adverse effects in order to propose reference levels, and a good description and structuration of the data from the first biological events to the observed adverse outcomes in order to characterize the mode of action of different pollutants. To do so, Adverse Outcome Pathways (AOPs), have been proposed by OECD few years ago as a conceptual framework that enables the structuration of the data and *in fine* a classification of pollutants according to the biological pathways they can trigger (2). In this context, data acquisition and structuration on chronic exposure to ionizing radiation (IR), naturally occurring but also reinforced due to anthropic activities, are of main interest. Indeed, IR share common early biological responses, namely oxidative stress induction via reactive oxygen species (ROS) production (3,4) or direct ionization of molecules (DNA, lipids, proteins, etc.), with numerous chemicals.

These primary processes can lead to adverse effects at larger scales i.e., individual and population via different pathways. In particular, reproduction, which directly impacts population dynamics, is the most radiosensitive individual parameter in many species, including invertebrates and more precisely *Caenorhabditis elegans* (5–9). However, the early molecular events of this reproduction decrease are still not all elucidated. To date, after chronic exposure to gamma rays during whole development of *C. elegans* (upon 50 mGy.h^-1^), previous results showed a decrease of brood size without any defect of hatching success (suggesting no embryonic damage at this range of dose rate with this exposure design) but concomitant with a decrease in sperm number and ovulation rate (10–12). In particular, we observed that ~90% of the brood size decrease can be attributed to sperm decrease, that can be possibly compensated by the increase of males in population (13), and ~10% to others, yet unexplained but probably linked to ovulation process (11).

Prior studies showed specific pathways in the germline involved in the response to chronic exposure to IR, in particular cell, stress response pathways (DAF-16/FOXO) and diverse effects on gametes proliferation (10,14,15). Among those pathways, the Ras/ERK (MPK-1) pathway is of particular interest because at the crossroad of observed effects. It acts to control several developmental processes in *C. elegans* hermaphrodite germline, including the regulation of spermatogenesis/oogenesis transition (16) (17) thus sperm count (18,19) and is also involved in stress response pathways, especially after irradiation (20,21). In *C. elegans*, MPK-1 expression is spatially and temporally dynamic, enabling to distinguish the various biological processes it regulates according to its zone of expression in the gonad (22). Notably, in the pachytene zone, MPK-1 controls the cell progression into meiosis (transition from distal to proximal pachytene) and the spermatogenesis/oogenesis transition, while in developing oocytes it controls oocyte maturation and ovulation. Apoptosis is another stress response pathway that was observed after chronic exposure in *C. elegans* (12,15) but its link with the modulation of the brood size remains to be demonstrated. While in *C. elegans*, prior studies under acute exposure to IR have shown that in male germ cells, neither DNA-damage induced nor physiological apoptosis occurs (23–25), the issue of induction after chronic exposure remains untreated. In response to DNA damages, the apoptotic pathway is triggered by other regulators than physiological apoptosis (i.e., normal apoptosis occurring during oogenesis) (25) but the downstream core apoptotic pathway remains the same, i.e. activation of the protease CED-3 caspase that cleaves specific substrates inducing killing and phagocytosis of the cell (26). In *C. elegans* germ cells apoptosis is restricted to late pachytene stage germ cells at the L4 stage (25).

Thus, in order to fill some knowledge gaps in the cascade of events leading to population decline after chronic exposure to ionizing radiation, we investigated the link between brood size decrease and pathways known to be involved in both stress response and developmental processes in the *C. elegans* germline, i.e., the Ras/ERK (MPK-1) pathway via dpMPK-1 staining in the germline, germ cell count, and the core apoptotic pathway via CED-3 loss of function mutant. To do so, nematodes were chronically irradiated at different life-stages as previously done (e.g., embryogenesis, early gonadogenesis, whole development) (11). In all scenarii, brood size, ovulation rate and sperm number were assessed.

## MATERIAL AND METHODS

### Strains and maintenance

The following strains were provided by the Caenorhabditis Genetics Centre (University of Minnesota, St. Paul, MN, USA): Bristol N2 strain was used as wild type, MT1522 (*ced-3(n717))* strain was used for the analysis of apoptotic pathway. Nematodes were maintained as described previously (11).

### Antibodies and reagents

The following antibodies and reagents were used in this study: Sigma monoclonal anti-activated MAP kinase (diphosphorylated ERK-1&2) antibody (M8159), Donkey anti-Mouse IgG (H+L) Highly Cross-Adsorbed Secondary Antibody, Alexa Fluor Plus 555 (InVitrogen, Waltham, MA, USA), Vectashield antifade mounting medium with DAPI (Vector Laboratories, Inc., Burlingame, CA, USA).

### Irradiation design

Worms were synchronized and irradiated in the Mini Irradiator for Radio Ecology (MIRE) facility at the IRSN laboratory, (Cadarache, France) as previously described (11). Wildtype worms (later stained with DAPI and MPK-1 antibody) were irradiated according to three scenarii of exposure (S1 Fig; SC1 – *in utero* until embryogenesis; SC2 – *in utero* until the beginning of gametogenesis; SC3 – *in utero* until the end of development). MT1522 strain was irradiated during whole development only (i.e., SC3).

### Sample preparation for dpMPK-1 assay and germ cell count

Following different exposure scenarii, wildtype N2 worms were all collected at the L4-YA (Young adult) stage, rinsed in M9 and quickly dissected and treated for dpMPK-1 (Sigma) staining according to Gervaise and Arur, 2016 protocol (22). MT1522 strain, ~20 worms per condition (CTRL vs SC3) were picked in individual petri dishes for reproduction assay, while the rest were rinsed in M9 and treated for DAPI staining and sperm counting.

### Reproduction Assay

Cumulated larvae number were quantified for each individual (20 per scenario) for eight days. Nematodes were transferred to fresh NGM petri dish every 24 h. In addition, newly laid eggs were quantified every 6 h after the transfer every day.

### Spermatids Quantification

Samples were washed three times with M9 and 15 to 25 worms per replicate were mounted on slides pre-coated with Poly-Lysine (10 μL at 0.1%). Worms were immobilized with 3 μL of 0.1 mM levamisole and dissected using a 0.45 μm gouge needle, then fixed with 2% paraformaldehyde. The freeze-cracking method was used to remove the cuticle (27). Slides were then fixed in methanol/acetone (1:1) for 20 min at −20 °C, then rinsed three times in 1x-PBST. Slides were stained with DAPI mounting medium for 2 h at +4 °C in a dark chamber. Images were obtained with Axioobserver ZEISS Z1 microscope (Carl Zeiss AG, Oberkochen, Germany) equipped with a DAPI filter system, at 40× and 12-bit resolution and Z-stacked for each spermathecal. Spermatids were quantified using the FIJI (Fiji Is Just) ImageJ 2.1.0/1.53c software (28).

### Image acquisition and quantification of dpMPK-1 intensity and sperm number

For dpMPK-1-stained wildtype worms, images were collected on a Zeiss LSM780 confocal microscope (Carl Zeiss, https://www.zeiss.com/) at the Zone d’Observation en Microscopie” (ZoOM) plateform at the “Commissariat à l’énergie atomique et aux énergies alternatives” (CEA, Cadarache, France) using a 40x objective (Plan Apo water N.A. 1,2). Images were acquired in 12 bits using Zen black software (SP2 v.11.0, 2012, Carl Zeiss). Alexa Fluor 555 conjugated to anti-mouse secondary antibodies (InVitrogen, Waltham, MA, USA) was excited with a 561nm Diode-Pumped Solid State (DPSS) Lasers and emitted light was collected from 564 to 573nm using the MBS 561 filter. DAPI was excited with a 405nm Lasers and emitted light was collected from 450 to 495nm using the MBS 405 filter. The measured intensity at different areas (zone 1: 80μm, and zone 2: −1 oocyte – see Figure 1) was weighted by the intensity at the loop area (0μm) chosen as the reference following the method of Eberhard et al (29) (see S2 Fig). The intensity is thus the ratio of the intensity at 80μm/0μm for zone 1, and −1 oocyte/0μm for zone 2. Cell proliferation on DAPI stained gonads was measured using the spots tool on IMARIS software v.9.9 (Oxford Instruments, Abingdon, UK).

**Fig 1.**
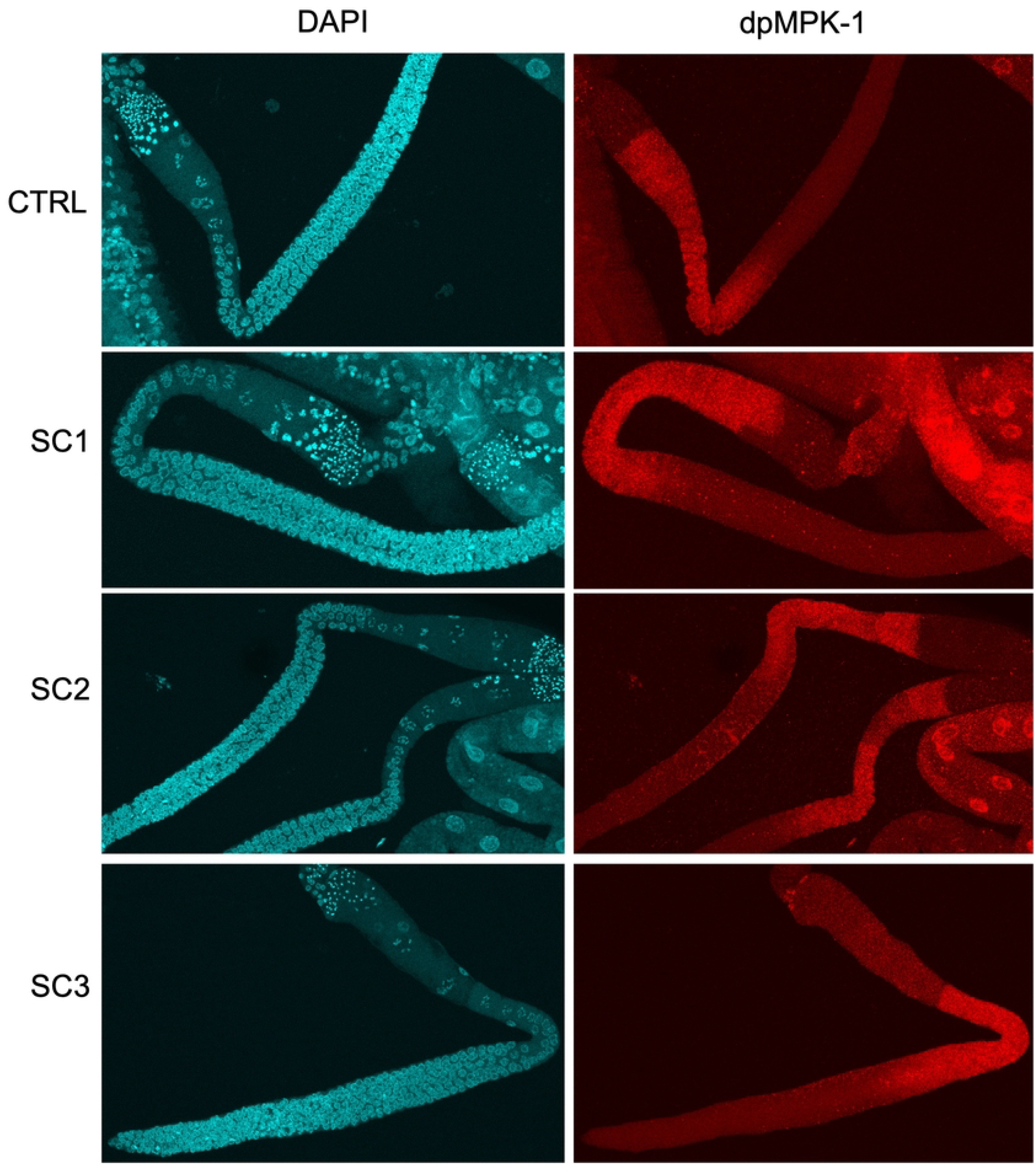
MPK-1 phosphorylation is induced in early pachytene (Zone 1) in wild-type germlines following irradiation at different life stages. (SC1: *in utero* until embryogenesis; SC2: *in utero* until the beginning of gametogenesis; SC3: *in utero* until L4/YA – whole development – see S1 Fig) ‘*’ p-value < 0.05

For MT1522 worms, images were acquired using ZEISS apotome as previously described (11). Sperm number was manually counted using the ImageJ/FIJI software (28).

### Statistical analysis

All data were obtained with a simple random sampling. Count data (spermatids count, total brood size, ovulation rate, mitotic cell count) were analysed as functions of irradiation conditions (i.e., SC1, SC2, SC3) using a GLM (General Linear Model) following a quasi-Poisson or gaussian distribution. For statistical significance of intensity, data was log-transformed to follow a normal distribution, means between conditions were compared using an Anova. For each test, post-hoc analysis was conducted (Dunnett for parametric tests) and p-values were adjusted using the Bonferroni correction. An alpha risk of 5% was taken as significant. Statistical analysis was conducted on R Studio software, Version 1.1.423 (© 2009–2018 RStudio, Inc., Boston, MA USA), using the following packages for statistical tests and data visualization: ‘ggplot2’, ‘ggpubr’, ‘dplyr’, ‘tidyr’, ‘car, ‘multcomp’.

## RESULTS

We investigated the involvement of two molecular pathways in the reprotoxic response to chronic exposure to ionizing radiations at different life stages in *C. elegans*: Ras/ERK (MPK-1) pathway and apoptotic pathway through CED-3. S1 Fig shows the experimental design and different scenarii of exposure seen in all the following figures.

### 1.1 MPK-1 is activated in the pachytene zone of the germline upon chronic exposure to IR

Upon exposure to many stressors, including IR, MPK-1 is activated, i.e., double-phosphorylated (dpMPK-1). In this study, the expression of dpMPK-1 in the germline was measured as described by (22). S2 Fig shows the different zones of expression of dpMPK-1 in *C. elegans* hermaphrodite germline and the areas where dpMPK-1 intensity was measured. Fig 1 shows examples of confocal microscopy images of dpMPK-1 and DAPI stained *C. elegans* L4/YA (young adult) hermaphrodite germlines after each exposure scenarii.

Fig 1 shows the intensity of dpMPK-1 expression in zone 1 and zone 2 as defined in S2 Fig. Important inter-individual variability of dpMPK-1 raw intensity was observed within conditions, especially for conditions with lower sample size (e.g., SC1) (S3 Fig). Therefore, to alleviate this variability, a reference zone in which no difference of intensity was observed between conditions was selected to weigh the raw intensity. No significant difference was observed between conditions in the loop area (S3 Fig), and intensities were similar to zone 2, therefore the loop area was selected as the reference zone. Then, the ratio of intensity between the zone of interest and the reference zone was used to compare conditions as show below (Fig 2).

**Fig 1.**
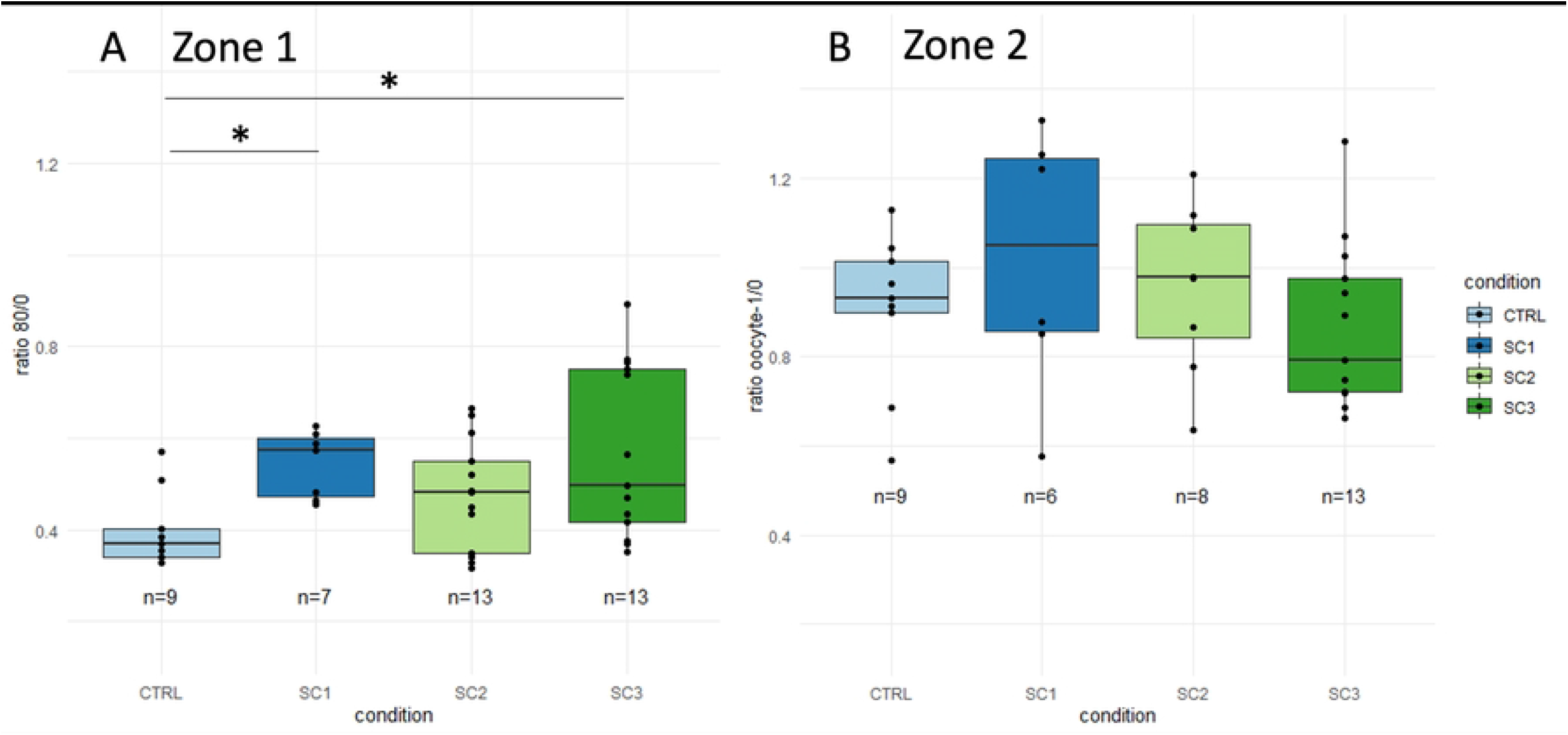
Analysis of dpMPK-1 spatial expression in N2 *C. elegans* hermaphrodite germline after chronic exposure to IR at different life-stages. (A) dpMPK-1 intensity in Zone 1 (intensity at 80μm weighted by intensity at the reference area 0μm – see S2 Fig) for each condition (SC1: *in utero* until embryogenesis; SC2: *in utero* until the beginning of gametogenesis; SC3: *in utero* until L4/YA – whole development – see S1 Fig) (B) dpMPK-1 intensity in Zone 2 (intensity at oocyte −1 weighted by intensity at the reference area 0μm) for each condition (Anova, Dunnett post hoc with Bonferroni adjustment, ‘*’ p-value < 0.05)

As shown in Fig 2, in zone 2, where dpMPK-1 controls the oocyte-linked processes (maturation, ovulation) (22), no significant difference between irradiated conditions and control was found. On the other hand, in zone 1, while the measurement of intensity at 40μm from the loop showed no significant difference between conditions (S3 Fig), dpMPK-1 is significantly over expressed at 80μm from the loop after irradiation throughout development (SC3) and after irradiation throughout embryogenesis (SC1). This can suggest a precocious activation of MPK-1 in the germline after irradiation, possibly affecting cell proliferation and meiotic progression, and later germ cell apoptosis. Considering this increased activation of MPK-1 in zone 1 and the possible link with cell proliferation and progression into meiosis (Section 1.2), we further investigated the effects of IR on cell proliferation in the mitotic zone of *C. elegans* gonad. In the same way the role of apoptosis in this process was assessed using *ced-3* mutants (Section 1.3). In both cases, a focus was made on SC3 condition which showed the highest increase of dpMPK-1 staining.

### 1.2 Total cell number and density in the mitotic zone of the gonad is unchanged upon chronic exposure to IR

Methodology for cell proliferation analysis in the mitotic zone is shown in S4 Fig. Fig 3A shows the mitotic zone length for each condition, while Fig 3B shows total number of cells per mitotic zone for each condition.

**Fig. 2.**
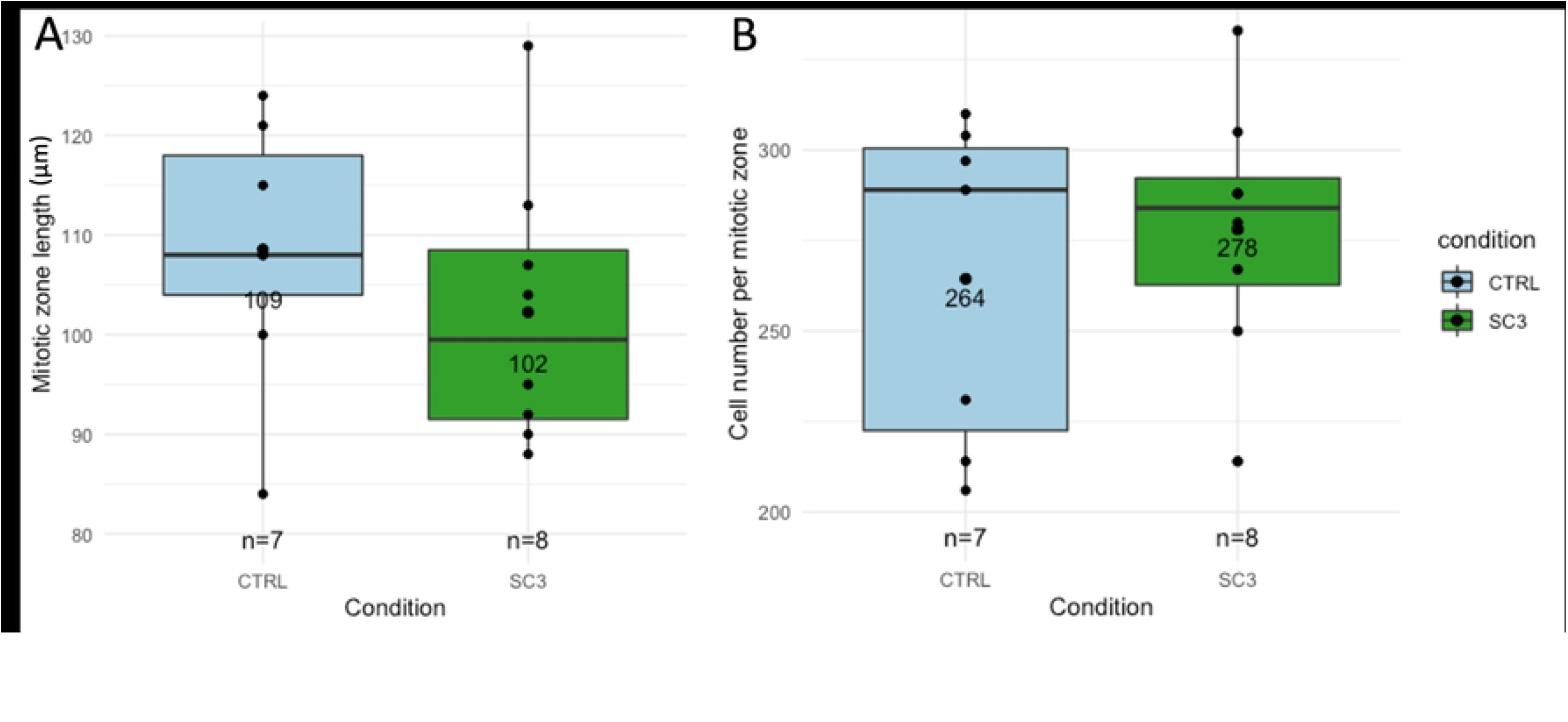
Analysis of cell proliferation in the mitotic zone of N2 *C. elegans* hermaphrodite germline. (A) Mitotic zone length for each condition (CTRL= 109±13 μm, SC3= 102±14 μm, GLM, Dunnett contrasts, p-value=0.37) (B) Cell number per mitotic zone (CTRL= 264±45, SC3= 278±36, GLM, Dunnett contrasts, p-value= 0.51)

A slight decrease of mitotic zone length can be observed in irradiated worms (Fig 3A; mean±SD for CTRL= 109±13 μm, SC3= 102±14 μm, GLM, Dunnett contrasts, p-value=0.37), however non-significant. Similarly, no difference was observed the total number of cells in irradiated worms compared to controls (Fig 3B; mean±SD for CTRL: 264±45, SC3: 278±36, GLM, Dunnett contrasts, p-value= 0.51). Therefore, it seems that IR has no effect on cell proliferation in the mitotic zone of the germline under our conditions.

### 1.3 Apoptotic pathway induces ovulation rate decrease but no effect on sperm upon chronic exposure to IR

Apoptotic loss of function *ced-3(n717)* worms were analysed under the same conditions as wildtypes N2 worms to understand the effects of apoptosis in the gamete response after irradiation. Fig 4 shows the total brood size, sperm count and ovulation rate of irradiated and controls *ced-3(n717)* (Fig 4 A,C,E) and N2 worms adapted from (11) (Fig 4 B,D,F).

**Fig 4.**
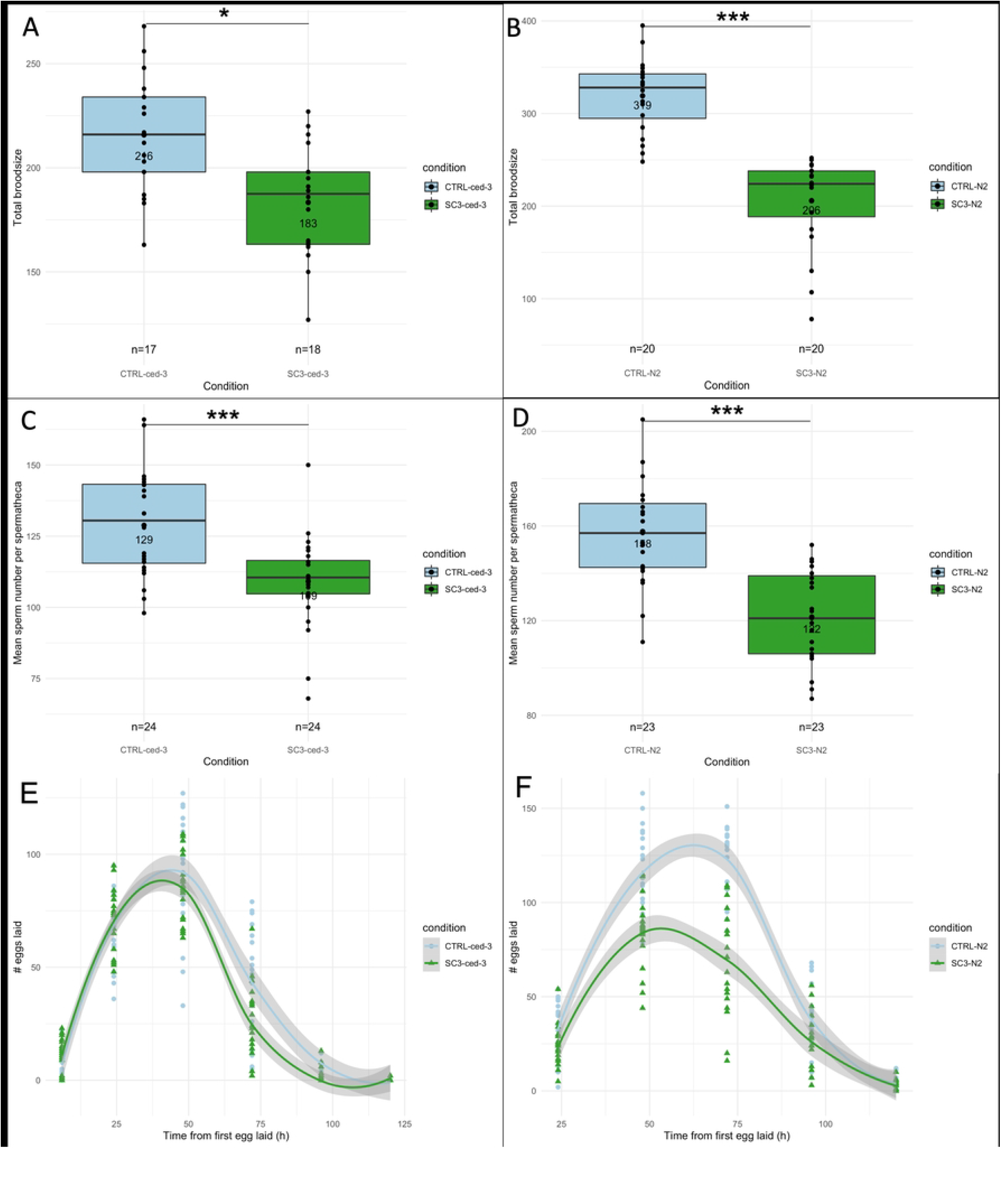
Analysis of radio-induced reprotoxic effects in non-apoptotic *ced-3(717)* mutant worms vs N2 worms. *ced-3(n717)* vs N2 analysis of brood size, sperm count and ovulation rate (A,C,E and B,D,F respectively) CTRL vs SC3: irradiated during whole development; GLM, Dunnett post-hoc with Bonferroni adjustment; ‘*’ p value < 0.05, ‘***’ p value < 0.001)

While brood size decrease is observed in irradiated *ced-3(n717)* worms compared to controls (Fig 4A, −15%, p-value = 0.02), a higher decrease is observed in N2 between irradiated worms and controls (Fig 4B, −35%; p-value < 0.001). Sperm number is reduced in irradiated *ced-3(n717)* worms compared to controls (Fig 4C, −15%; p-value < 0.01) equivalent to the *ced-3(n717)* brood size decrease observed in Fig 4A. A similar decrease in sperm number was observed in N2 under the same conditions of irradiation (Fig 4D) (−23%; p-value < 0.001). On the contrary, no effect of radiations on ovulation rate in irradiated *ced-3(n717)* worms was observed (Fig 4E), contrarily to what we observed in N2 (Fig 4F, −35%, p-value < 0.001). Thus, irradiated non-apoptotic mutant worms show a similar reduction of sperm count and brood size decrease with N2 worms, but no effect on ovulation rate contrary to N2.

## DISCUSSION

### 3.1 MPK-1 activation in the pachytene region reveals involvement of apoptosis more than defects in cell proliferation

Our study aimed to decipher the underlaying mechanisms leading to the adverse outcome “brood size decrease”, crucial for population dynamic, in order to link the different biological levels of this radio-induced reprotoxic effects. Based on results from previous studies (10–12,15,25,30), we focused on processes involved in germline development and stress response such as the Ras/ERK (MPK-1) pathway and apoptotic core machinery. As previously described, MPK-1 is involved in many processes such as the regulation of sperm count (19,31), germ cell proliferation (32–34) and differentiation (35–38), but also in stress response pathways (20,21,25,39). No significant difference in dpMPK1 expression was observed in the ‘ovulation zone’ (i.e., zone 2), meaning that maturation of oocytes and ovulation processes are not affected by IR under our condition of exposure. Thus, the 10% ovulation rate decrease that was previously observed (11) might be the consequence of prior effects occurring during oocyte proliferation and differentiation. Similarly, we observed no significant difference of dpMPK-1 expression in the loop area (cells transitioning from late pachytene to early diplotene). On the contrary, our results show an overexpression of dpMPK-1 upon exposure throughout the whole development in the early to late pachytene zone (i.e., zone 1), where it controls the progression of cells into meiosis, and apoptosis as previously described (21,29). In addition, it is also concordant with the overexpression of *egl-1* observed in similar conditions since its activation is required for IR–induced *egl-1* transcription (15,21). MPK-1 activation in this zone can be involved in several processes, notably sperm-promotion and cell proliferation (40–43). In our conditions, it is not linked to sperm promotion since a decrease of sperm count was observed. In addition, germ cell proliferation investigation in the mitotic zone did not show any significant result neither on mitotic zone length nor on germ cell density in our conditions, with a mean number concordant with the literature (i.e., ~236 mitotic cells) (44). To conclude, the MPK-1 pathway is activated upon chronic exposure to IR in our conditions only in the pachytene region of the gonad showing a possible induction of the apoptotic pathway.

### 3.2 Apoptosis is responsible for oocyte decrease but not spermatocyte decrease

Previous studies had shown the induction of apoptosis in *C. elegans* via several pathways upon chronic exposure to IR. Notably, the expression of the proapoptotic gene *egl-1* (15) was increased as well as the presence of apoptotic cell corpses (10), and differential regulation of miRNA family *mir-35*, also involved in fecundity and embryonic development (12,43). Despite this upstream evidence, it was not clearly known to which extent the apoptotic pathway contributes to downstream brood size decrease. Our analysis of the core apoptotic pathway revealed that radio induced apoptosis is not involved in sperm decrease but is involved in ovulation rate decrease. Indeed, sperm number is decreased after irradiation in *ced-3(n717)* mutants in the same way as in N2 (11), which means that this decrease occurs via another mechanism than apoptosis. On the other hand, the ovulation rate is not affected by irradiation in *ced-3(n717)* mutants, contrary to what was observed in N2, which means apoptosis is involved in the decrease of the ovulation rate in N2 after chronic irradiation, probably even independently from the regulation by spermatozoa, notably via the Major Sperm Protein (18,45). In conclusion, brood size decrease in *ced-3(n717)* mutants can be attributed to sperm decrease only. This confirms that the brood size decrease in wildtypes is not only due to sperm decrease but also to ovulation decrease, via the apoptotic pathway.

From our study, we suggest that two mechanisms act in parallel that affect spermatogenesis and oogenesis independently both contributing to brood size decrease (Fig 5). The causal mechanisms of sperm decrease remain unknown, but we show that neither the Ras/ERK (MPK-1) pathway nor apoptotic pathway are involved in this decrease.

**Fig 5.**
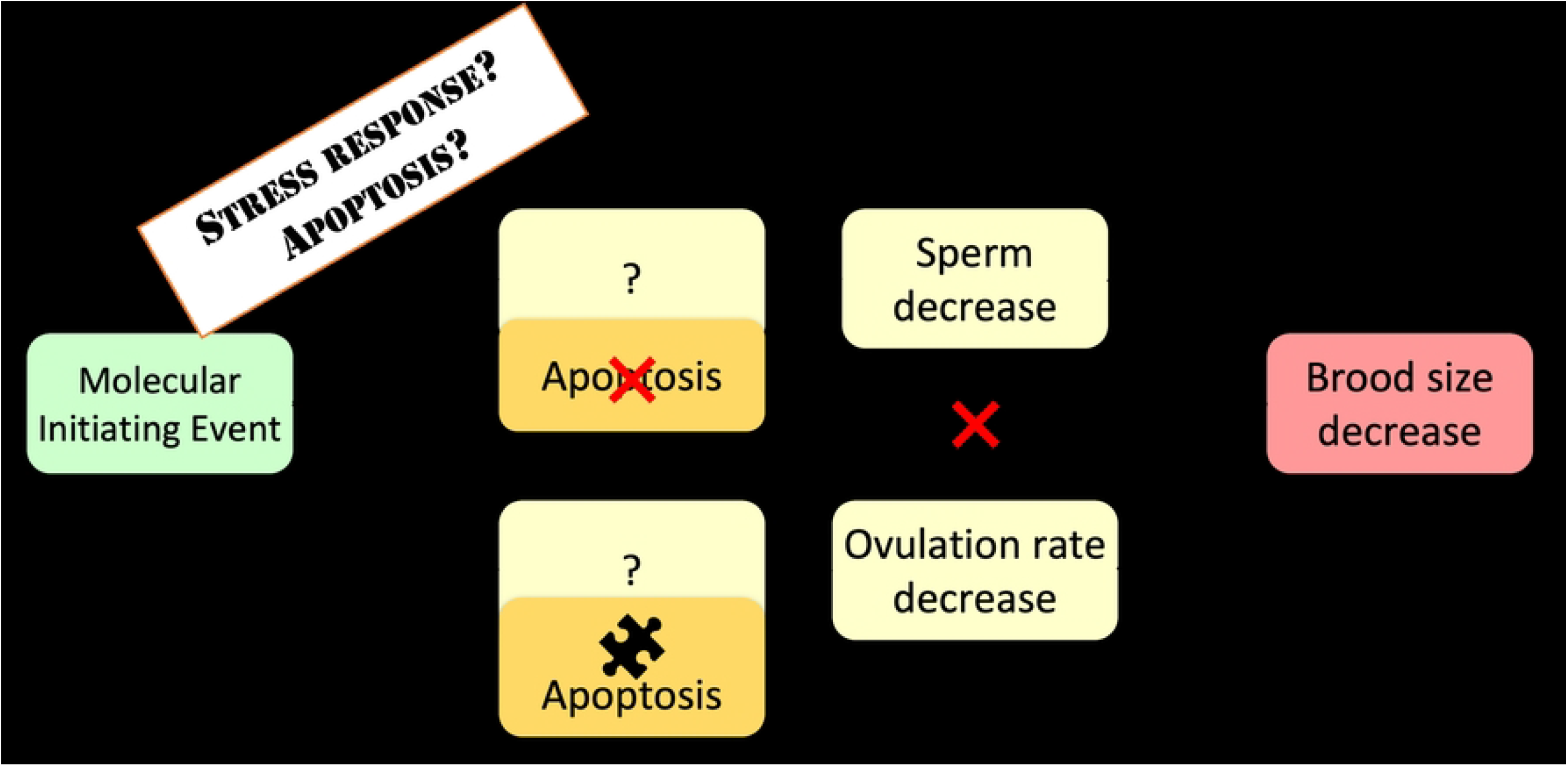
Proposed steps leading to of brood size decrease Adverse Outcome in hermaphrodite *C. elegans* exposed to chronic ionizing radiations. Two mechanisms act in parallel that affect oogenesis and spermatogenesis independently and lead to brood size decrease. Oocytes are reduced directly and only via apoptosis, while spermatocytes are reduced via another yet unknown mechanism. Question marks represents missing key events or key events relationships. Puzzle symbol represents key events that were validated in this study. Red cross represents key events and key events relationships that were invalidated in this study.

### 3.3 Causes of sperm decrease: perspectives from our results and the literature

In previous studies, sperm count was shown to be decreased only after exposure throughout the whole development, and not only in precocious stages (11) (with the same dose rate) nor throughout gametogenesis ((10) with an equivalent total dose). These results suggest that there is cumulative damage during larval development, culminating in sperm decrease at the adulthood. In many species, sperm decrease is mediated via DNA damage-induced apoptosis (46–48). In the present study, it was shown that apoptosis is not involved in sperm decrease in *C. elegans*. Similarly, no effect were observed in the length and the cell density of the mitotic zone, showing that sperm count decrease is apparently not linked to germ cell proliferation defect. However, several previous studies on *C. elegans* identified a potential slowdown of germ cell proliferation showing increase of cell cycle arrest in the mitotic zone of the gonad after acute exposure to IR (25), or chronic exposure from 37 mGy.h^-1^ (15) or a decrease in expression of proteins linked to the MCM complex involved in DNA replication and critical for cell division (14). Therefore, a decrease in germ cells proliferation speed probably due to DNA damage and repair could directly induce a decrease in the number of spermatozoa since they are produced on a finite time-lapse. Considering all these lines of evidence, this hypothesis cannot be discarded and needs to be further investigated to fill the knowledge gap and finally identify the missing key event of brood size decrease in *C. elegans*.

## CONCLUSION

Following our previous studies showing a decrease of hermaphrodite *C. elegans* reproduction upon chronic exposure to IR, this study aimed to investigate more specific mechanisms linked to gamete development and stress response. We showed effects in both oocytes and spermatocytes but via different mechanisms. Indeed, oocytes seemed to be only affected by apoptosis via the Ras/ERK (MPK-1) pathway which led to a slight decrease of ovulation rate. On the other hand, spermatocytes were not affected by apoptosis, but we still observed a reduction of sperm count that can be hypothetically attributed to a slowdown of proliferation in germ stem cells. Interestingly, this work can contribute to understand why after multigeneration exposure, it is observed that individual brood size still decreases continuously over time, despite an increase of males in the population to rescue fecundity (13). This continuous decrease could be partly explained by the 10% oogenesis decrease due to apoptosis throughout adulthood. Thus, only looking at the individual scale could enable to think that sperm decrease is critical to reproduction decrease (90%) contrary to oocytes (10%). However, on a larger scale, sperm decrease is rescued by the increase of males in populations, while oocytes keep decreasing and become mainly responsible for reproduction decrease at the population level. Our present result can be linked to effects at larger biological scales, giving a meaningful interpretation for ecological risk assessment. Further research is needed to determine the mechanisms of sperm decrease, possibly via cell proliferation defects and to understand the causes of radiosensitivity of these gametes in *C. elegans* and potentially other organisms.

## AKNOWLEDGEMENTS

The authors thank the IRSN for their financial support. All nematode strains were provided by the CGC, which is funded by NIH Office of Research Infrastructure Programs (P40 OD010440). We kindly thank Nicolas Dubourg for the dosimetry calculation. Support for the microscopy equipment was provided by the Région Provence Alpes Côte d’Azur, the Conseil General of Bouches du Rhône, the French Ministry of Research, the CNRS and the Commissariat à l’Energie Atomique et aux Energies Alternatives.

## SUPPORTING INFORMATION

**S1 Fig.**
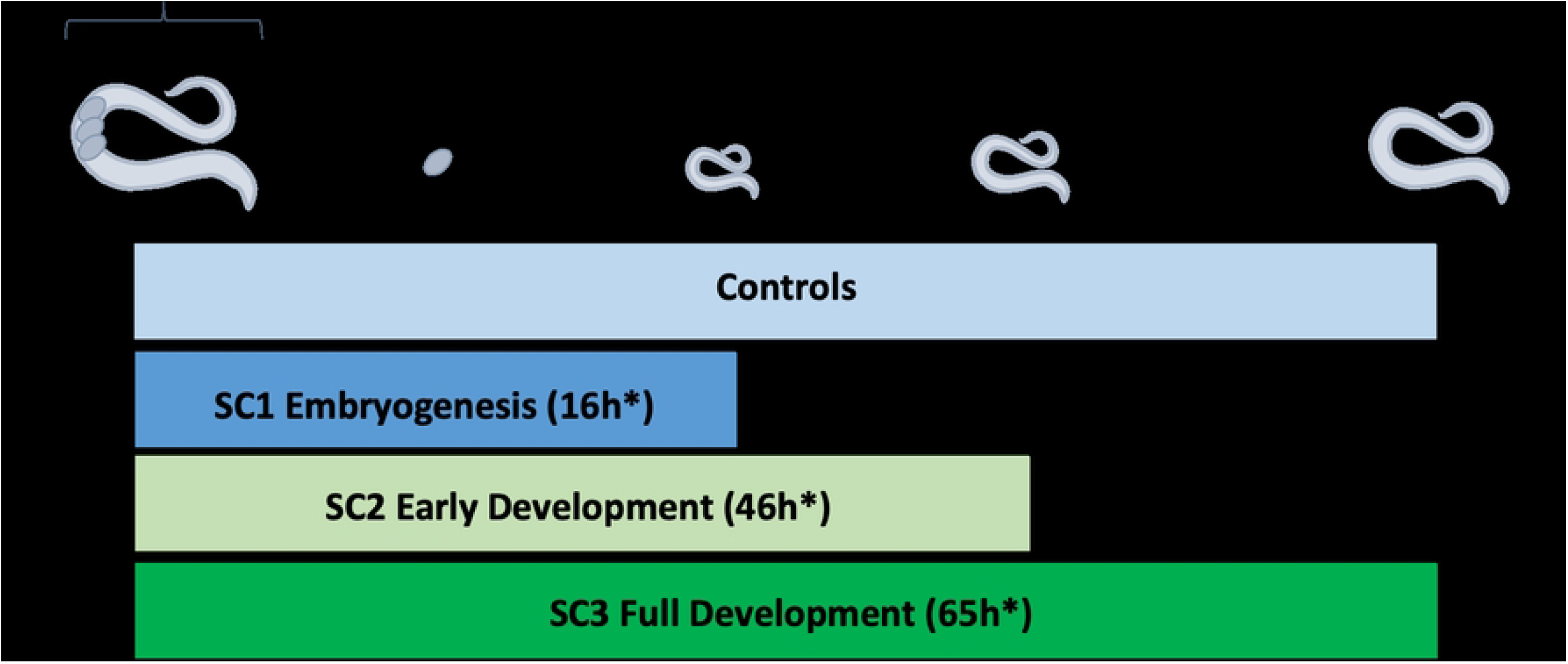
Experimental design for analysis of life stage dependent reprotoxic effects. (adapted from Dufourcq-Sekatcheff *et al* 2021). (SC: scenario, OP50: E. Coli strain, L1-L4: C. elegans larval stages, YA: Young Adult).

**S2 Fig.**
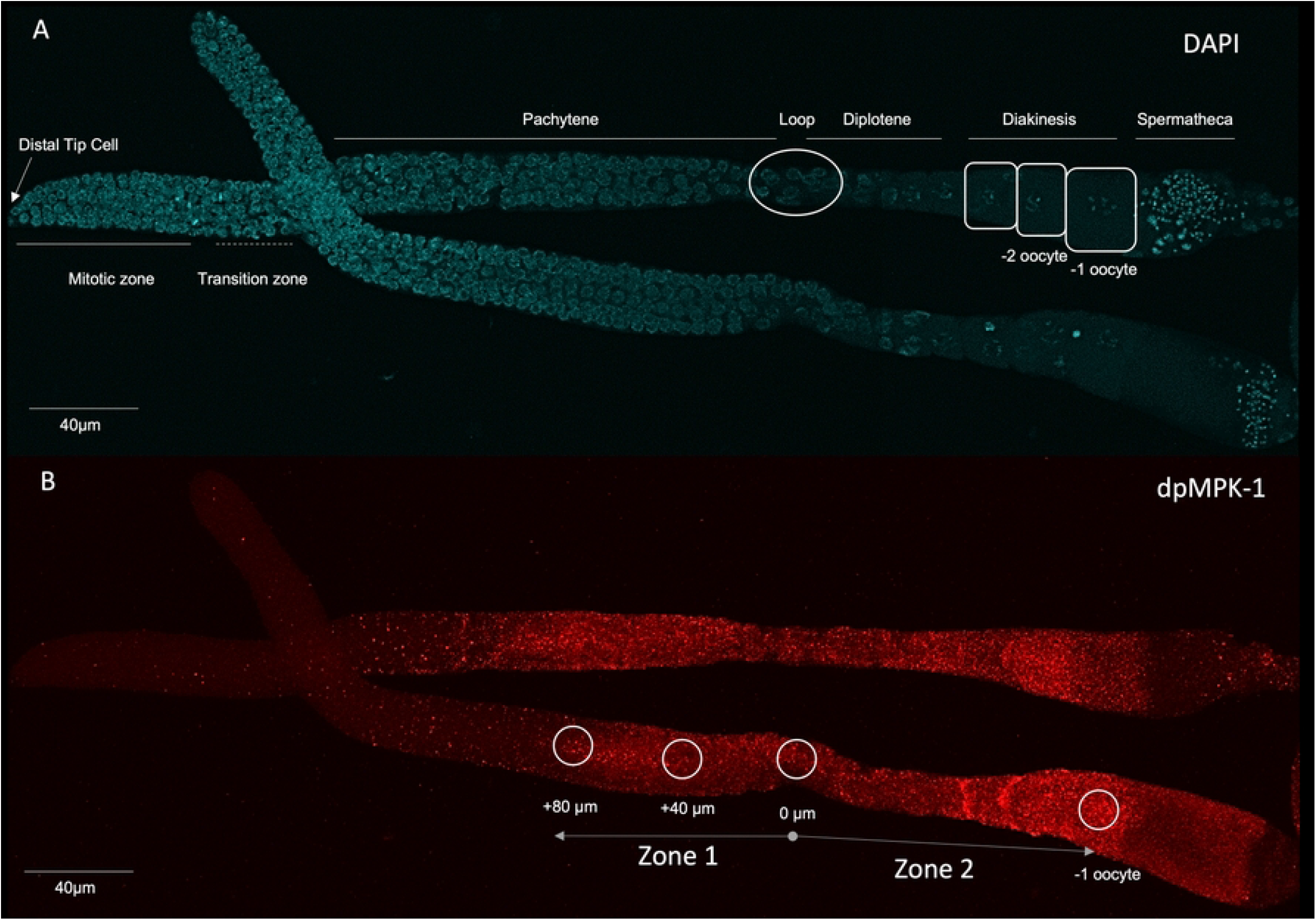
(A) DAPI-stained L4/YA hermaphrodite *C. elegans* germline. Germ cells differentiate from the distal end of the gonad (distal tip cell on the left) to the proximal end (spermatheca on the right). (B) dpMPK-1 expression in C. elegans dissected gonad (Zone 1 corresponds to expression in the pachytene stage, 2 measures of intensity were taken at 80μm and 40μm from the loop (0μm); Zone 2 corresponds to expression in the proximal oocytes).

**S3 Fig.**
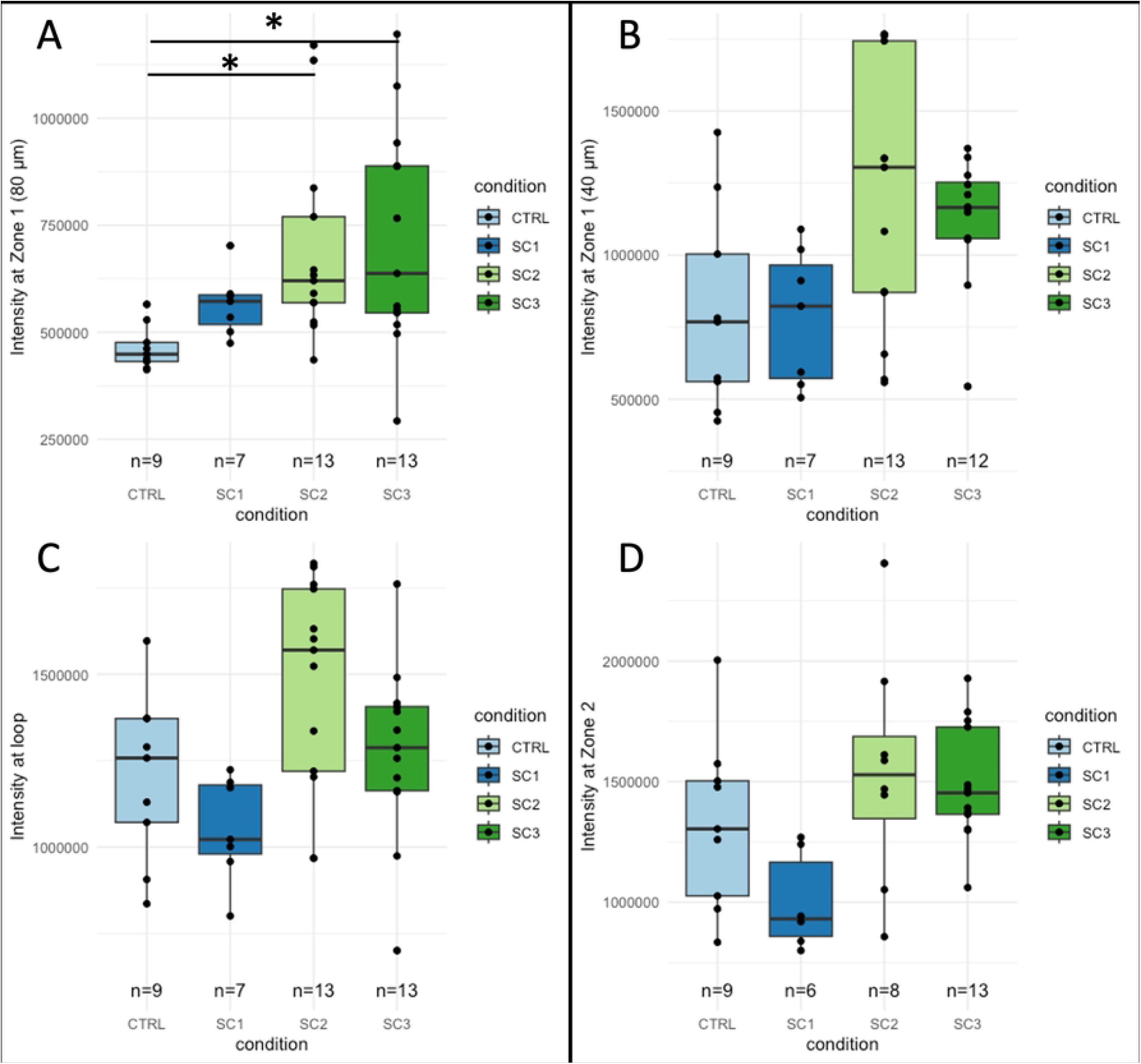
Raw intensity of dpMPK-1 in different areas of the germline for each condition of exposure. (A) Intensity in Zone 1 at 80μm from the loop area towards the distal end of the gonad (B) Intensity in Zone 1 at 40μm from the loop area towards the distal end of the gonad (C) Intensity in the loop area (D) Intensity in Zone 2 (−1 oocyte). (Kruskal Wallis, Dunn test with Holm adjustment, ‘*’ p-value < 0.05)

**S4 Fig.**
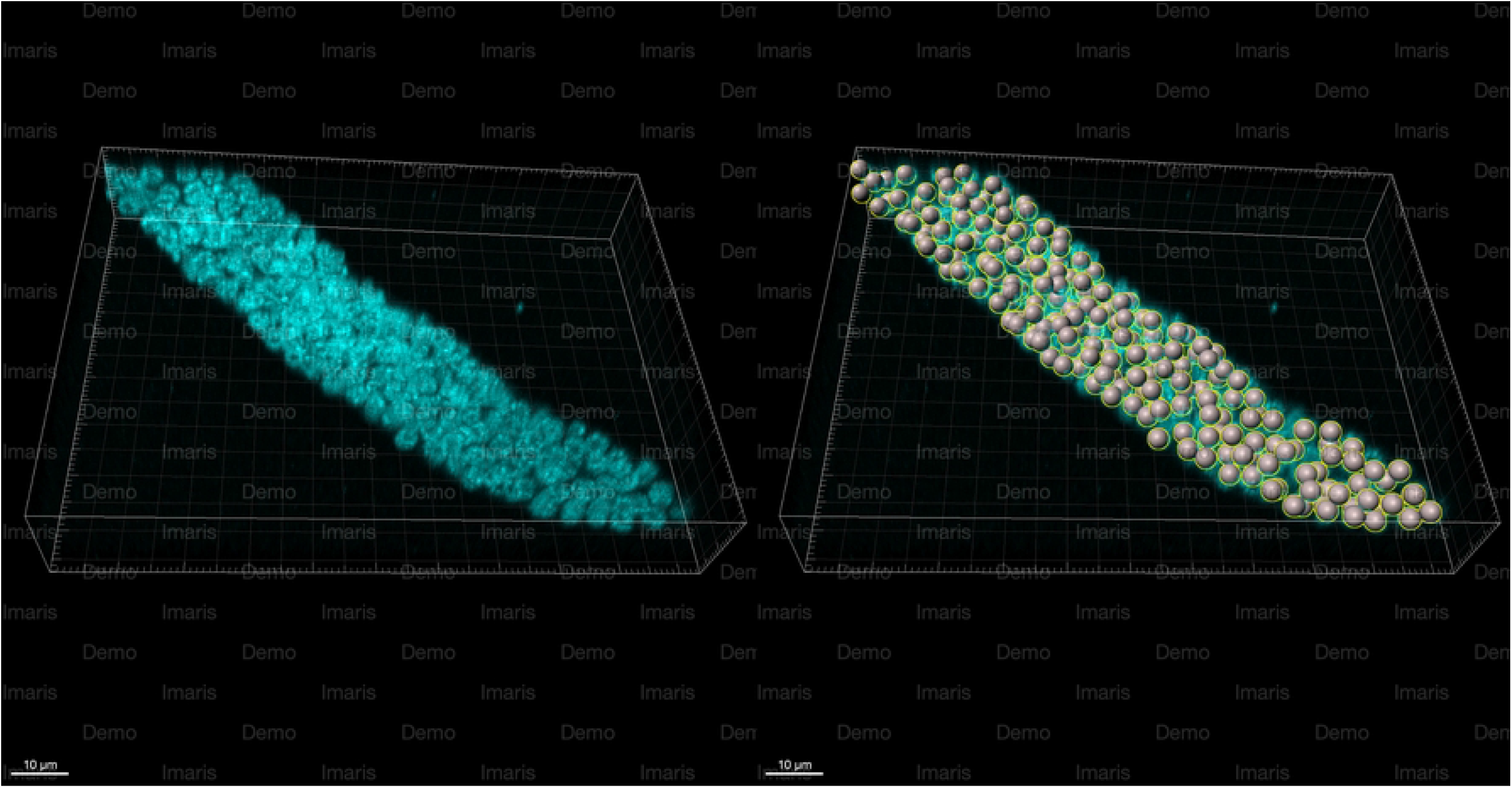
Automated cell counting of mitotic cells in the *C. elegans* distal gonad. Mitotic Zone extends from Distal Tip Cell to first meiotic cells. Left: 3D z-stack of a DAPI-stained dissected gonad, right: automated spot counting of the same gonad using IMARIS software.

## Notes

### Competing Interest Statement

The authors have declared no competing interest.

